# Development of a Time-based Drug Screening Platform for Improved Clinical Translation

**DOI:** 10.1101/2025.10.24.684432

**Authors:** Elizabeth A. Hughes, Lindsay L. Davenport, Andrew Woodward, Shravan Patel, Jamie Lee, Christopher Zdyrski, Aleksandra Pawlak, Karin Allenspach, Jonathan P. Mochel, Eugene. F. Douglass

## Abstract

In 2022, the average cost for a single drug’s development increased by 15%, reaching $2.3 billion. In drug development, oncology has the highest attrition rate with 95% of new drugs failing Phase 2 clinical trials alone. Consequently, innovative approaches are needed to assess clinical utility prior to proceeding with human trials. Current drug development follows the theory that “Best Potency = Best Drug” which is a concentration-centric paradigm focused on optimizing drug-target interaction (K_d_) to improve the concentration of drug necessary to elicit its therapeutic effect (**IC**_**50**_). However, both *drug concentration* and *exposure time* contribute to clinical efficacy, yet most laboratory studies focus on **IC50** alone. This manuscript characterizes drug potency and kinetics of ten oncology drugs in three different cancer cell lines and integrates in vivo pharmacokinetic-pharmacodynamic models to identify how these parameters relate to real-world clinical outcomes. Our analyses revealed that C_max_ normalization is necessary for concentration response data as IC_50_ alone does not account for drugs that are studied at sub-therapeutic concentrations. Additionally, the temporal effect of drug efficacy varies between cell lines and dose, where some drugs are unable to overcome the proliferation rate of the cells to induce a decrease in disease progression. This work aims to enhance the design and implementation of drug regimens by understanding the time-dependence of clinical efficacy and cytotoxicity.

## Introduction

### Translational Challenges

In drug development, the failure rate in drug translation is 90%, with the highest attrition rate observed in oncology therapies.^1,2^ In 2022, the cost to bring a single drug to market increased to $2.3 billion.^34,5^ Drug development begins with the identification of a drug target or phenotypic screens, where the first two steps focus on concentration-response characterization (Fig. 1, black). Based on the crystal structure, high-throughput drug screening, and/or *in silico* modeling, a series of compounds is developed and optimized based on drug-binding affinity (Kd), the midpoint of binding affinity in concentration-response curves with protein assays (Fig. 1A).^6^ Then, cell-based assays are used for determining efficacy, which establishes the concentrations at which off-target effects in normal cells and drug potency in disease model cell lines are achieved (Fig. 1B, left). The threshold concentrations of off-target toxicity and drug efficacy are often defined by the 50% inhibitory concentration (IC50), and their ratio defines the therapeutic range.^7,8^ These early concentration-response curves are critical for designing initial dose-escalation studies in animal studies and later phase 1 clinical trials (highlighted in yellow in Fig. 1B).

**Figure 1.**
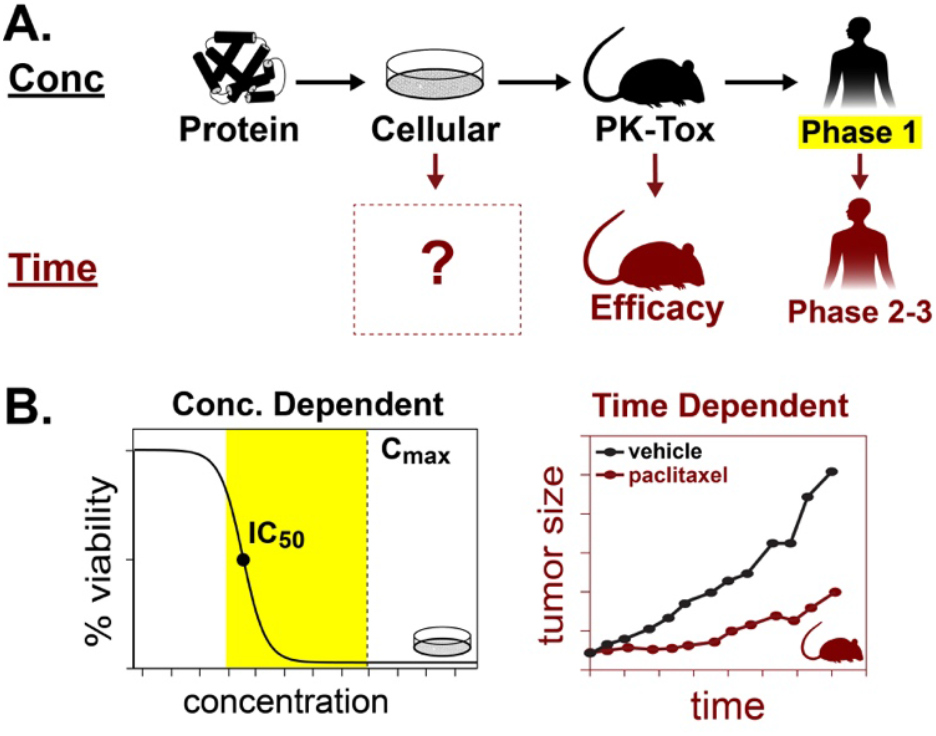
A. Conceptual Overview of Drug Development. B. Drug discovery begins with concentration–response studies in protein and cell assays, which guide PK–toxicity testing in animals and humans through dose–response relationships. In contrast, in vivo efficacy is typically assessed as a time– response, creating a knowledge gap since drug kinetics are often not examined until Phase 2 trials, contributing to late-stage attrition.

After cell line validation, the top compounds are tested in animal models for initial pharmacokinetic studies and drug efficacy studies. ^7–9^ Critically, this step makes a transition from concentration- to time-response efficacy metrics (Fig. 1B). The step from cell-based models to animal models is a major bottleneck in the drug development pipeline, where only 30-60% of drugs proceed to Phase 1 trials due to a lack of efficacy seen in animal studies.^2,8^ Phase 1 trials confirm the maximum plasma concentration (C_max_) that can be administered to human patients and to determine the optimal regimen for drug efficacy studies.^2^ After the ideal dosing regimen(s) are determined, the drug efficacy is tested in Phase 2-3 clinical trials, which is the next major bottleneck, primarily due to a lack of efficacy (Fig. 1A, red). The in vitro effectiveness described is not always predictive of *in vivo* effectiveness, due to differences in drug distribution and exposure time. These in vivo effects are modeled using pharmacokinetic-pharmacodynamic (PK-PD) models, which incorporate both drug potency and kinetic rate constants, e.g., k_dmg_ and k_kill_ for drug-induced signaling and cytotoxicity, respectively. We believe that one of the most significant limitations is the lack of understanding regarding the kinetic parameters of these drugs prior to animal studies (Fig. 1A, red box). To bridge the gap between *in vitro* efficacy and clinical translation, innovative approaches are needed to analyze the clinical utility of drugs in cell-based assays.^3^

We are not the first to propose kinetic characterization of drugs at the in vitro stage. In 2016, Sorger and colleagues introduced an important extension of traditional dose–response assays (24–72 h endpoints) by incorporating cell density measurements at both the beginning and end of experiments to capture growth-rate effects. The “Sorger-GR” model represented a major advance because it aligned in vitro drug-efficacy metrics with clinical response categories of progressive disease (growth rate >1), stable disease (growth rate =1), and tumor regression (growth rate <1). However, Sorger-GR does not capture two key kinetic parameters routinely used *in vivo*, as in the Simeoni-TGI model: k_dmg_, the immediate downstream signaling rate constant, and k_kill_, the rate constant of the terminal cytotoxic step.

In 2004, Simeoni and colleagues developed a pharmacodynamic-pharmacokinetic model to fit drug response in mouse xenograft studies and the rate of proliferating cells within the tumor in both treated and untreated models (k_pro_).^10,11^ Simeoni highlights 4 key steps in drug response: the first is that the drug must damage the cells, followed by downstream killing.^10,11^ The downstream killing is divided into three steps to account for the observed lag in the response, similar to the downstream effects following initial target binding. While only using three killing phases or steps is an oversimplification of drug response due to variations in mechanisms of action across drug classes, this phenomenological term can be applied to multiple oncology drugs to recapitulate drug response. A critical feature of Simeoni-TGI is its prediction of transient viability peaks within time–response curves defined by 1/k_dmg_. Here, we evaluate the validity of applying the Simeoni-TGI framework to *in vitro* time–response data, extracting k_pro_, k_dmg_ and k_kill_, and relating them to clinical response. By doing so, we demonstrate that kinetic parameters offer predictive power comparable to traditional potency metrics, thereby establishing a conceptual bridge between *in vitro* pharmacology and real-world clinical efficacy. ^10,11^

## Results

### Study Design

To understand how well concentration and kinetic metrics outlined in Fig. 2 correlate with clinical responses, an assay optimized for continuous monitoring of cell viability was chosen. This also enables integration into the existing infrastructure of drug screening centers. The most commonly used *in vitro* viability assays are traditionally endpoint kits, such as Presto Blue, Alamar Blue, Cell Titer Glo, and MTT assays. While many new methods have been developed to monitor drug responses over time, the Promega Real-Time Glo (RTG) assay was used due to its fidelity with traditional endpoint assays and compatibility with plate readers and robotics used in drug screening centers.^14^

**Figure 2.**
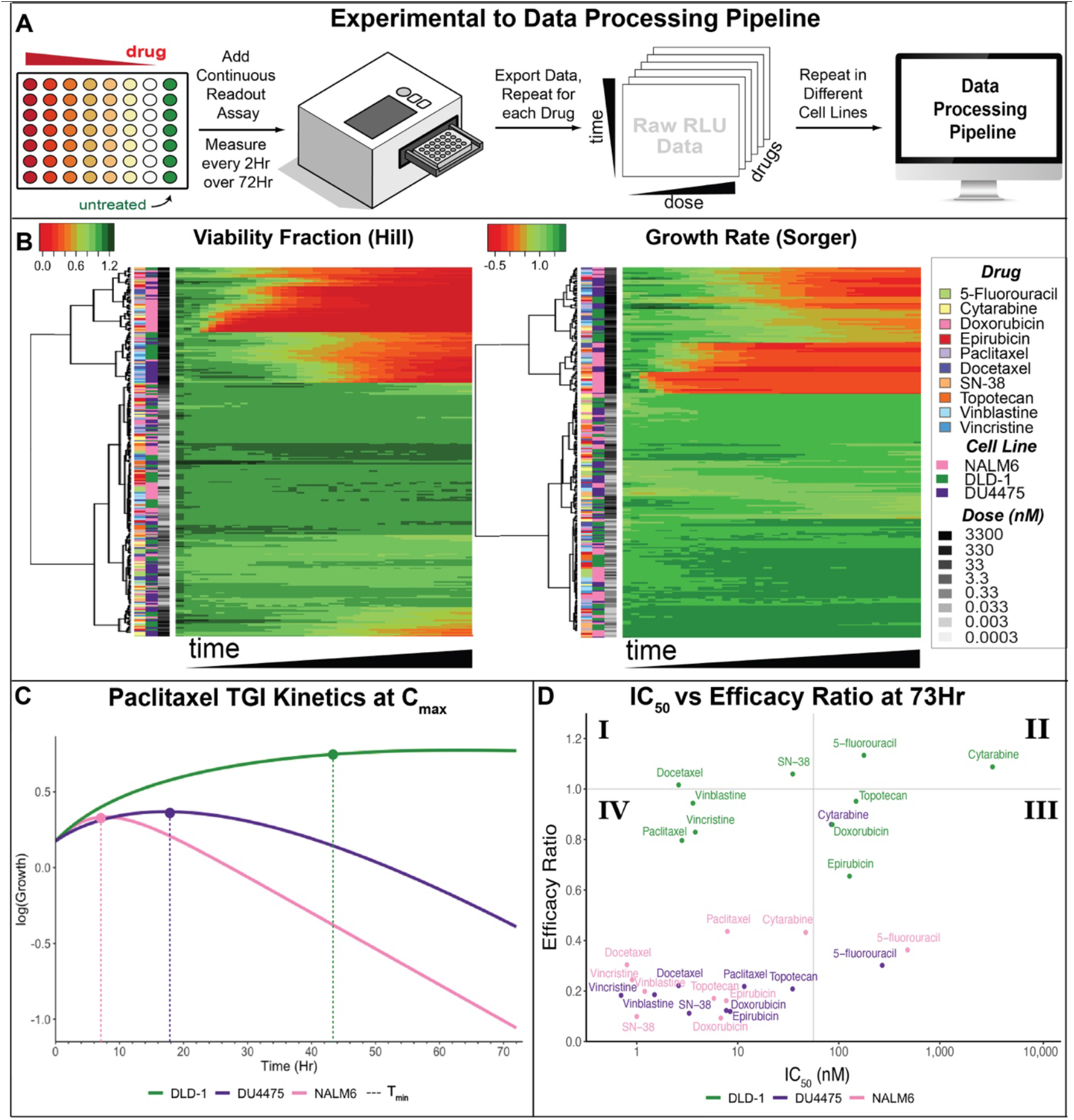
Data Generation and Analysis Overview. **A.** Generation of experimental data through the proposed Data Analysis Pipeline. Two drugs from each of the top 5 oncology drug classes were chosen and tested across 3 cell lines. **B**. Heatmap overview of all experimental data acquired by both Viability Fraction (left) and Growth Rate (right). In each heatmap scale, **red** is associated with dead cells and **green** with alive cells. The rows are color coded from left to right drug, cell line, and dose tested (nM) according to the legend (far right), while the x axis is each of the 36 time points (1Hr to 72Hr) **C**. An example of the TGI Kinetic Model of Paclitaxel at close to the C_max_ (Conc used is 50 nM, clinical C_max_ is 100 nM) was plotted in log scale. The arrows denote the peak of the simulated curve showing how long the drug needs to be in the system to induce cytotoxicity (where the **k**_**DMG**_ > **k**_**PRO**_). **D**. A comparison of IC_50_ in log10 scale versus Efficacy Ratio (ER) in log2 scale for all drugs and cell lines. The Efficacy Ratio is the **k**_**PRO**_ divided by the **k**_**DMG**_. The horizontal line denotes where ER = 0. The vertical line denotes the average **IC**_**50**_ value across the data.

Three cell lines with varying growth rates and sensitivity to oncology drugs in our focused chemotherapy library were used. These cell lines included leukemia (the most chemosensitive), triple-negative breast cancer (moderately chemosensitive), and colorectal cancer (chemo-insensitive/chemoresistant), and were chosen for their prevalence of clinical data and ranking among the top 10 most common cancer types by incidence. DLD-1 (colon cancer) has a doubling time of 20-33 Hrs^15–17^, NALM6 (leukemia) of 19-24 Hrs^17,18^, and DU4475 (breast cancer) of 70-90 Hrs^17,19^. Two chemotherapy drugs were selected from the standard drug classes used in typical treatment regimens for these cancers. Colon cancer is often treated with 5-fluorouracil or Irinotecan (the prodrug of SN-38), typically in combination with a platinum drug.^20^ The treatment regimen for Triple Negative Breast Cancer can include 5-fluorouracil, Doxorubicin, Epirubicin, Paclitaxel, Docetaxel, Vinblastine, or a combination of one or more of these drugs.^20^ Leukemia can be treated with Doxorubicin, Cytarabine, or Vincristine given as a monotherapy or in combination.^20^ Topotecan was included to have a second drug with the same mechanism of action as SN-38. (Fig. 2A).

### Design of Computational pipeline (Figure 2A)

The raw RLU data were processed according to traditional drug efficacy models —Viability Fraction (Fig. 2B, left) and Growth Rate (Fig. 2B, right)^21,22^ — as outlined in the methods section, and the results were presented as heatmaps in RStudio. Both heatmaps use red as dead cells/killing and green as living cells/proliferation. Visualizations of the concentration-response and growth curves for all drug-cell line pairs can be found at https://lizahughes.shinyapps.io/RTG_App/. The Simeoni TGI Model was applied to the Raw RLU data using the Ordinary Differential Equations (ODEs) from the original publication.^10,11^ The parameters were fit and used to calculate the maximum Efficacy Ratio by dividing the max k_PRO_ by the max k_DMG_ for each drug-cell line combination (y-axis of Fig. 2C) and the Concentration Threshold at 73 Hours (Table 1). The Concentration Threshold for each time-point is also reported in the Supplemental Data. The raw data and code used to generate the figures can be found at https://github.com/fingolfn/rtg_simeoni.

### Viability Fraction: Endpoint “Snapshot” of viable cancer cells

Viability fraction (VF) is a traditional data normalization method used to fit concentration-response models, such as the Hill Equation. This metric calculates the number of viable cells in the presence of the drug divided by the vehicle control. As the most common efficacy metric used, we first evaluated how this metric changes over time (Fig. 2B columns) and how it is affected by drug-type, drug-concentration, and cancer model (Fig. 2B rows)

While traditional viability fraction metrics provide an endpoint “snapshot” of the number of viable cancer cells relative to our untreated control, when monitoring the process over time, it was revealed that a minimum time is required to induce cytotoxicity. There are two main groups separated in our heatmap: responders (top cluster) and non-responders (bottom cluster). Concentration was found to be 90% of the driving factor between the two major clusters (Supp Fig. 2A). Once killing occurs (concentration > IC_50_), drug mechanism and cancer context are the primary factors in killing speed. Within our responding group (Supp. Fig. 2C heatmap), a transition peak was consistently observed, which captured the transition from proliferative to toxic cellular states. This peak captures the minimum time required for cytotoxicity (t_min_) and is not directly captured by traditional endpoint measurements in Viability Fraction or Growth Rate frameworks (which assume monotonicity).

There are three main clusters of responders, separated by the speed of killing: fast, medium, and slow (left Fig. 2B, Supp. Fig. 2C). Principal component analysis identified the driving factors of killing speed. The fast-killing cluster is the group of cell line-drug pairs that induce cytotoxicity before 24 hours. Fast killing is dominated by the cell line NALM6 and topoisomerase inhibitors (Anthracyclines and topoisomerase 1 inhibitors) (left side for PC1 in Supp Fig. 2C). The tubulin inhibitors (Taxanes and Vinca Alkaloids) are only fast in NALM6. Whereas the topoisomerase inhibitors are the only drug class that is fast acting in all three cell lines, and no other drug class is fast in DLD-1 or DU4475 (right side of PC1 in Supp. Fig. 2C). This indicates that fast responders may be driven by mechanisms and cancer type rather than proliferation speed. Medium-speed killing is typically between 24-48 hours, which is dominated by tubulin inhibitors in DLD-1 and DU4475. In DU4475, in addition to the tubulin inhibitors, a response to lower concentrations of Epirubicin and Doxorubicin was observed, whereas in DLD-1, responses were observed to 5-fluorouracil, SN-38, and Doxorubicin. There is also the presence of 5-fluorouracil and lower concentrations of Docetaxel in NALM6. The final cluster is the slow-killing group, where toxicity is not observed until 48-72 hours (bottom rows of the heatmap in Supp Fig. 2C). This is dominated by Cytarabine, 5-Fluorouracil, and Docetaxel in DU4475, and by Epirubicin and Topotecan in DLD-1. Overall, the drug’s mechanism of action is the primary factor in killing speed, but cell line and, potentially, cancer type also have clear effects on killing speed. This analysis highlighted that different “HIT” compounds would be identified depending on the time point (24, 48, or 72 hours).

### Growth Rate: Percentage of Viable Cells compared to treatment start

Growth rate (GR) is the proposed summary kinetic metric used in the Sorger-GR model, which is calculated from the change in cell number over time. Comparison of GR and VF changes over time similarly shows that both metrics are highly sensitive to the timepoint at which they are measured, suggesting a complexity more consistent with in vivo PK-PD models such as Simenoi-TGI (Fig 2B).

Comparison of VF and GR metrics across our dataset revealed similar concentration-trends to Viability Fraction and the conserved presence of a “transition peak” that affects drug-killing speed. While Sorger and colleagues propose using two time points for practical reasons, this approach would also miss the minimum time of drug exposure required to slow proliferation (t_min_), as indicated by the gradual shift to red in Fig. 2B (right). Similarly to Viability Fraction, there is a divide between non-responders and responders driven by concentration (Supplementary Fig. 3B, left), but compared to Viability Fraction, there is less killing (red). Another key difference is that in the responding group, only fast and medium-speed responders are present, with a limited number of slow responders.

**Figure 3.**
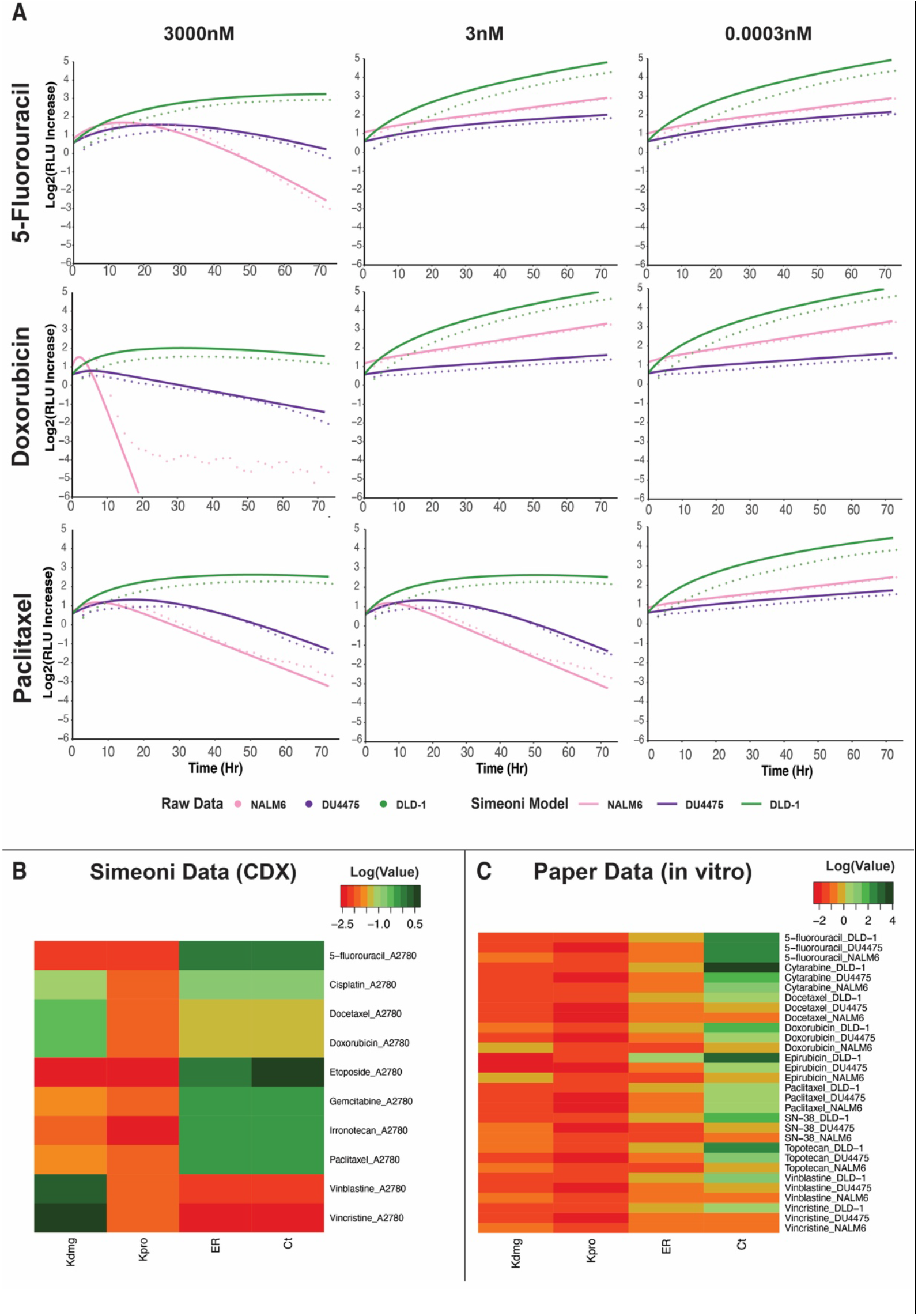
Validation of The Simeoni-TGI Model in *in vitro* Data. **A.** The fit of the TGI Model across three different drug classes and concentrations (solid line) overlaid with the average of triplicate replicates (dotted line) **B**. A heatmap overview of the experimental results of Simeoni’s of various drug treatments in a cancer derived xenograft (CDX) of A2780 (ovarian cancer) in mice **C**. The results of using a derivation of Simeoni’s PK-PD model in our experimental data.

The fast-killing cluster in growth rate is associated with cancer- and drug type by NALM6 and topoisomerase inhibitors, where DU4475 and DLD-1 only have a fast response to the topoisomerase inhibitors (left side of PC1 in Supp Fig. 2D). The main difference is that in DLD-1’s Growth Rate, the only fast responders seen are the Anthracyclines. In DLD-1, the topoisomerase 1 inhibitors are unable to inhibit growth rapidly, despite a significant decrease in viability fraction. Similarly, tubulin inhibitors and Cytarabine remain fast in NALM6, whereas 5-fluorouracil is no longer identified as a fast-growing agent. In medium responders, a strong emphasis on DU4475 and DLD-1 was observed, with a higher presence of NALM6 than in the viability fraction. In NALM6, lower concentrations of tubulin inhibitors and Doxorubicin were observed, compared with a high concentration of 5-fluorouracil. In DLD-1, responses are observed at high concentrations of topoisomerase inhibitors, tubulin inhibitors, and 5-fluorouracil. As with the viability fraction, a minimum concentration is required to induce an effect; however, the Growth Rate data show that the cell line and the drug’s mechanism of action determine the rate. Overall, the trends are similar to those seen in the viability fraction matrix; however, the Growth Rate may reveal specific mechanisms of action and kinetic information. The main difference between these two metrics is that viability fraction is a snapshot of a drug’s ability to kill cancer cells, whereas Sorger’s Growth Rate is a continuous measurement of the drug’s ability to inhibit cancer growth.

### Comparison of VF and GR

Viability Fraction metrics emphasize a snapshot of how well a drug can kill cancer cells compared to the control, while Growth Rate metrics highlight the kinetic nature of a drug’s ability to inhibit cancer cell proliferation. Critically, neither the Viability Fraction nor the Growth Rate framework is explicitly fit to the drug-efficacy speed limit (T_min_), as reflected in the transition peaks across datasets. Therefore, the Simenoni TGI model was applied, revealing novel clinical insights.

### Efficacy Ratio/Concentration Threshold

While Sorger-GR’s GR provides a better overall comparison of disease progression than Viability Fraction, the model cannot recapitulate the complexity of drug kinetics. When applying Sorger-GR to our *in-vitro* methods, the predicted curves are monotonic and do not capture the complexity of drug kinetics (Fig. 2C/Supp. Fig. 3). Sorger’s model, while easier to apply, makes it harder to capture and identify the transition between proliferation and cytostatic/cytotoxicity we see in the continuous readout data (T_min_). Next, Simeoni’s fitting procedure was applied to *in vitro* assays and found that these *in vivo* models can successfully recapitulate *in vitro* responses (Fig. 3C/Fig. 4). Critically, the summary statistics provided direct insights into drug speed, which neither VF nor GR directly addresses. For each drug-cancer pair, the k_PRO_, k_DMG_, and k_kill_ were determined for each concentration (Fig. 4C/Supp Table 1). K_dmg_ and k_kill_ have been previously defined, while K_pro_ defines the basal proliferation rate and is closely related to the control GR measured with the Sorger-GR model.

**Figure 4.**
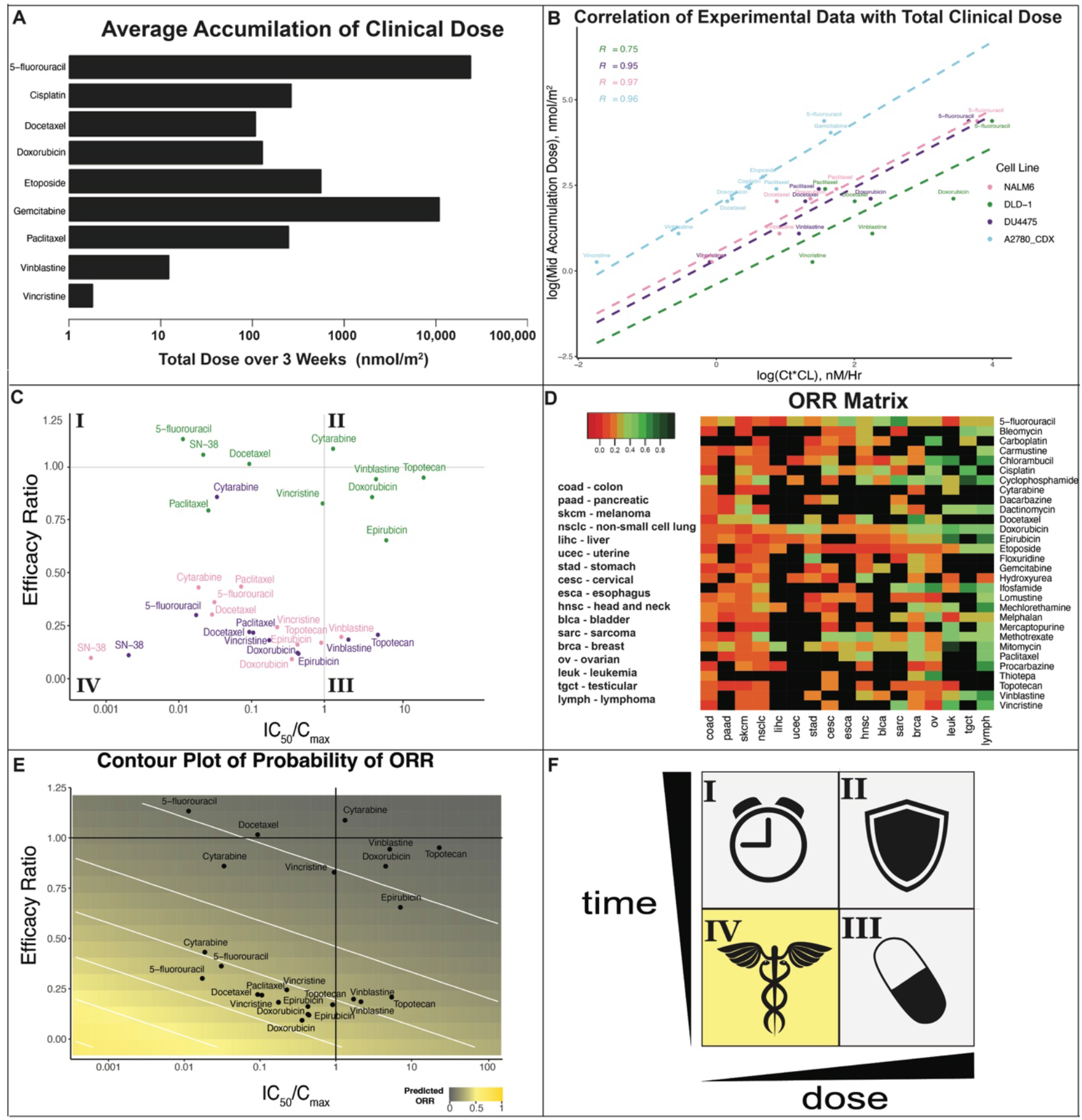
Clinical Comparison and Development of Potential Translation Criteria. **A.** A Bar plot showing the average accumulation dose over 3 weeks of treatment according to the standard regimens as outlined in Simeoni’s publication. **B**. The correlation between the experimental data and the average clinical accumulation dose over three weeks compared to the product of Concentration Threshold (Ct) multiplied by the Clearance in Human for that particular drug. This is used to evaluate how well Concentration Threshold from the Simeoni Fit can predict the clinical dose given to patients. For DLD-1 the R value was 0.75, but for all other cell lines the R value is above 0.95. **C**. A comparison of IC_50_/C_max_ versus Efficacy Ratio (ER) scale for all drugs and cell lines. This plot is an expansion of Fig. 2D to determine how clinical concentrations shift drug efficacy. **D**. Patient Overall Response Rate (ORR) for phase 2 clinical trials encoded by the NCI and literature curation of monotherapy response of 30 drugs in 17 cancer types. In this heatmap, **red** is associated with poor patient response and **green** with good patient response. **E**. A contour plot of probability of patient response rate developed using a generalized additive model using the data from Fig. 4D and Fig. 4C. **F**. Prediction of Clinical Indication where clinical efficacy can fall into four different quadrants, indicating as a function of either dose or time selectivity.

Critically, Simeoni and colleagues defined a summary kinetic-efficacy term, the threshold concentration for tumor eradication (Ct), that reconciles all kinetic and concentration-summary statistics into a single aggregate term. Defined as the steady-state concentration required to reduce tumor burden completely 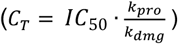 The C_t_ can be determined by the efficacy ratio (ER), the ratio of the maximum k_PRO_ to the maximum k_DMG_, multiplied by the IC50. (See reformulation of Simenoi in the methods). ER provides a metric for assessing the competition between the cell’s ability to proliferate and the drug’s cytotoxicity. This metric can be used to see differences between drug classes and cell lines, as highlighted in Fig. 3B. For an ER greater than 1, a steady-state concentration higher than the IC_50_ would be required to slow cancer cell growth *in vitro*, and an ER at 1 (denoted by the grey horizontal line in Fig. 3D) would indicate that the steady-state concentration at the IC_50_ would induce stabilization of cancer-cell number (analogous to clinical “stable disease”). Whereas an ER lower than one, a steady-state concentration lower than the IC50 would be sufficient to induce cytotoxicity. While most cancer chemotherapies are not administered for 73 hours at a single steady-state concentration due to their toxic nature,^20^ this summary statistic provides a way to combine drug kinetics and concentration sensitivity into a more effective metric for ranking drugs for clinical translation.

Drug response can be separated into four main quadrants, based on their concentration dependency (x-axis) and their kinetic dependency (y-axis) in Fig. 3D: I (top left): High potency, slow effect; II (top right) low potency, slow effect; III (bottom right) low potency, fast effect and finally IV (bottom left) high potency, fast effect. Using Simeoni’s Parameters to quantify the Efficacy Ratio as a function of the cell line’s proliferation rate relative to the drug’s damaging rate, and to compare it to traditional concentration metrics such as IC_50_, yielded similar trends in the heatmaps, presented in a concise format. Drugs such as 5-fluorouracil had an IC_50_ of ∼500nM in all three cell lines, but their kinetic selectivity varied between cell lines; the ranking of drug damaging rate from slow to fast is DLD-1, NALM6, and DU4475. While 5-fluorouracil has poor dose selectivity, its typical clinical exposure is very high with a C_max_ of ∼16,000 nM. The other example is Paclitaxel, which shows a much lower IC_50_ (∼7 nM), but It takes much longer to work in DLD-1 than in NALM6 and DU4475, similar to 5-fluorouracil. From the chart, it was observed that DU4475 is the fastest responder and most sensitive to Paclitaxel, which aligns with the fact that the Taxanes are standard in TNBC regimens, while at lower concentrations (30 nM) NALM6 responds quicker than DU4475. Even though the Taxanes are not standard of care for leukemias, this could indicate a potential area for repurposing or a better understanding of clinical data and pharmacokinetics.

### Application of the Simeoni Data

The Simeoni-TGI Model was applied to the raw data using the set of ordinary differential equations as outlined in the original publications and Eqs 1-2 in the Methods section.^10,11^ The parameters estimated from the fit were then used to create simulated data to determine the goodness of the fit and any deviations from the data (Fig. 3A). Three concentrations (the highest, median, and lowest) and three example drugs with different mechanisms of actions were chosen to highlight the ability for Simeoni’s ODEs to recapitulate the patterns from the experimental data (Fig. 4A). Overall, the *in vitro* data had an overall higher damaging rates than the xenograft study; this could be due to tissue and/or drug exposure or other pharmacokinetic parameters (Supp Table 1,3).

As mentioned earlier, the initial doses chosen for phase 1 clinical (dose-escalation) studies are first evaluated in preclinical mouse studies. Although the dosing regimens can vary across cancer type and patients, Simeoni wanted to see how well predicted concentration thresholds (Ct) compared to total active dose in human patients. To estimate that accumulation dose was taken over three weeks of treatment (Fig. 4A) and compared to the Ct multiplied by the Clearance (Supplemental Table 1,3) to see if there was a correlation between the values. Experimental values from Simeoni’s publication were extracted to compare their analysis to our method. SN-38 and Irinotecan were excluded from the prediction of active dose due to the inability for most cell lines to convert irinotecan into the active SN-38 form, and since SN-38 is rarely given as is in the clinic (Fig 4B). To determine how well the *in vitro* system can predict active doses, we replicated this analysis for our data with Simeoni’s data in blue (R = 0.96). Our R values were similar to Simeoni’s, NALM6: 0.97 and DU4475: 0.95, except for DLD-1, which had an R value of 0.75. This may be because colon cancer regimens tend to be higher since it is generally resistant to treatment.^20^ Overall, it is encouraging that a high correlation between the *in vitro* data and the predicted active dose was seen.

One major limitation, as mentioned earlier, is that many commonly used drugs are highly effective despite their poor potency, such as 5-fluorouracil. To account for this limitation, the IC_50_ was divided by the average C_max_ (Supp Table 2) to create a new summary plot (Fig. 4C). When normalizing to C_max_, we begin to see a stronger clinical correlation with a typical drug according to dose alone. For colon cancer, 5-fluorouracil and SN-38 move further into Quadrant I to become some of the most potent drugs compared to the clinical dose. For TNBC, 5-fluorouracil moves from Quadrant III to Quadrant IV which is consistent with its role as the “backbone” of most clinical regimens for colon cancer.

To directly benchmark these in vitro metrics vs clinical efficacy, a dataset of monotherapy human clinical response data for 30 drugs in 17 different cancer lineages was compiled to create a matrix of cancer response (Fig. 5D). We believe by combining kinetic and dose selectivity metrics, an improved go/no-go criteria for clinical translation can be created (Fig. 5F). A generalized additive model was developed to evaluate the relationship of IC50/Cmax to the ER of a particular drug-cell line pair, to the corresponding drug-cancer ORR (Fig. 5E). When comparing to only using IC50 (Supp Fig. 2) there isn’t a significant increase in predictive power (change in Bayesian Information Criterion ∼1), but the main driver of ORR becomes the efficacy ratio of the drug rather than its clinical potency. Overall, IC_50_/C_max_ and ER metrics showed similar predictive power in related in vitro and clinical efficacy metrics.

## Discussion

### Context-specificity of drug action (cancer-type and/or patient-specific)

Although both Viability Fraction (Hill) and Growth Rate (Sorger) capture different information, when used together, we can better understand the “snapshot” of drug response (Viability Fraction) as well as the “kinetic” response of a drug (Growth Rate). Because Viability Fraction is a snapshot metric, cytotoxicity in *in vitro* assays can be observed while having an increase in cell count compared to before treatment. This is because tumor size and cell count depend on how rapidly cells are proliferating, and drug efficacy is a competition between proliferating cells and drug-induced cytotoxicity. Therefore, different drug “HITS” are identified when looking at concentration response versus growth inhibition.

While 72 hours is the time point typically used to report IC_50_ or GR, we can see that cytotoxicity is transient even at a single concentration (Fig. 3B, left). While Sorger proposes using two time points, when multiple time points are present, it can be seen that there is a minimum time of drug exposure to slow proliferation (Fig. 3B right). As a result, Sorger’s growth rate shows less top drug-cancer pains than viability fraction or IC_50_. When continuously observing these metrics over 72 hours (as opposed to a single time point), both metrics can highlight differences between fast- and slow-acting drugs, but they also reveal distinct factors that contribute to a drug’s clinical efficacy. One example is that 5-Fluorouracil can induce cytotoxicity quickly in NALM6 (less than 24 hours), but doesn’t slow growth until closer to 48 hours. Similarly, Paclitaxel (Fig. 3C) shows a strong difference in kinetic speed across cell lines. In NALM6, we see both a decrease in Viability Fraction and Growth Rate quickly; however, in DU4475 and DLD-1, we see a slower decrease in Viability Fraction, but we see a slowing in cancer growth between 24-48 hours. These results highlight how a single time point can lead to a misunderstanding of a drug’s translational potential, which may explain the poor efficacy observed in animal xenograft studies and clinical trials following *in vitro* studies. An additional factor Sorger mentions is that proliferation rates can vary across studies, and the IC_50_ changes over time, so determining the optimal single time point for drug efficacy may differ across studies and cell lines. Additionally, comparing the drug-cancer rankings over time highlights that some drugs may induce cytotoxicity relative to the control but are not effective at inhibiting cancer growth. This phenomenon occurs because a drug must act faster than the cancer’s ability to proliferate. Overall, we believe this indicates the need for improved cell models to understand drug potency alongside drug kinetics.

The goal was to determine whether Simeoni’s analysis could be taken a step further to assess whether cell-based assays could predict the patient’s overall response rate, creating a go/no-go model for precision medicine and drug development efforts. Where the theory is that there are four contributing factors in drug efficacy, and where it is located shows the driving factor contributing to lack of sensitivity: I: Poor Kinetic Selectivity, II: Chemoresistance, III: Poor Potency, and finally IV: Strong candidates for clinical translation. In Quadrant I are examples of drug-cell line pairs that would be highlighted as highly potent in drug screening data, given their low IC50, but the rate of damage is slower than the cells’ proliferation rate. These drugs take longer to induce the desired effect but are highly potent. This is a prime example of how concentration-dependent metrics might yield “false positives” at later time points, even when cell density is higher than at the beginning of the assay. Given that the rate of damage is slower than the rate of proliferation, we see low overall efficacy in Sorger’s Growth Rate. For example, in DLD-1, Docetaxel, the Growth Rate never gets below 0, showing that we never achieve cytostatic behavior (Supplemental Data). In Quadrant II, we see poor potency as well as poor kinetic selectivity. These drugs would most likely be “true negatives” in high-throughput screening. In DLD-1 at 73 hours, the maximum concentration (3000 nM) induced only a 35% Cytotoxicity with 5-fluorouracil and a Growth Rate of 0.24. Quadrant III includes drugs with a high IC_50_, but they are fast-acting drugs in the particular cell line. These are drugs that might not pass high-throughput screening due to their poor concentration sensitivity but could induce the desired effect rapidly. Again, we have 5-Fluorouracil, this time in NALM6 and DU4475. As highlighted above, 5-fluorouracil can rapidly induce cytotoxicity in NALM6 but only causes a slight delay in growth inhibition. However, at 3000 nM, a Growth Rate of -0.1 is observed at 73 hours (Supplemental Data), indicating a substantial decrease in cell density. Finally, we have Quadrant IV; these drugs are highly potent, fast-acting drugs. These drugs are examples of what might be “true positives,” in which we observe a strong response at lower concentrations that decrease cell density. An example is Paclitaxel in NALM6, where we observe a rapid switch from cell proliferation to death relative to the other cell lines (Fig. 3C) and an IC_50_ of 7.9 ± 0.1 nM (Supplementary Table 1). This model was then used to create a predictive model of patient response *in vitro* (Fig. 5E). Encouragingly, the model supports our general hypothesis that, as potency relative to the clinical dose and kinetic selectivity increase, patients are more likely to respond to a drug. While this model is limited to one cell line per cancer type, we believe it reinforces the idea that drug potency is important but has limited clinical translation potential, as both the clinical dose and drug kinetics play a strong role in driving clinical effectiveness.

Overall, it is understood that a drug’s efficacy depends on cancer type and patient-specific factors within each cancer type. These trends are conserved in our cell-based assays. Hematopoietic cancers are fundamentally different from solid tumors and are generally considered easier to treat and more “curable” – as evidenced by the clear separation between NALM6 and our solid tumor-derived cell lines. While triple-negative breast cancer tends to be one of the most chemosensitive of the breast cancer subtypes, there is still a strong preference for different oncology drugs, with the most commonly used being the taxanes, anthracyclines, and 5-fluorouracil.^23,24^ DU4475 has the clearest separation of drugs based on the mechanism of action; this may be because DU4475 is slow-growing and a more resistant line of triple-negative breast cancer. Of the three cancer types, colon cancer is typically the most difficult to treat, and we see that consistent with the lower number of drug “HIT”s seen in our screen. While colon cancer is typically more chemoresistant, DLD-1 is also a faster-growing cell line, suggesting that drugs with the highest efficacy must act more quickly to outpace its proliferation rate.

While cell line and, presumably, cancer type strongly influenced the ranking of the drugs, the mechanism of action also strongly influenced the speed of response for both viability fraction and growth rate (Supp Fig 1). Topoisomerase inhibitors were generally faster, especially at higher doses. These different killing rates across doses might be due to the multiple mechanisms of action within topoisomerase inhibitors, especially the anthracyclines, and to the fact that topoisomerase is highly expressed in the S phase of cell division.^13,25^ Tubulin inhibitors and antimetabolites are more dependent on the cell cycle, generally take 24-48 hours to act, and are expressed in the G_2_/M phase.^13,25,26^ Additionally, these drug classes showed less concentration dependency, indicating a cleaner mechanism of action.^27–29^

### Limitations in Concentration-only metrics

While traditional concentration metrics provide essential information about drug efficacy, they are limited in clinical translation because they focus solely on potency rather than the drug’s kinetic ability to slow or reverse tumor growth. Additionally, pharmacokinetics need to be considered since many effective drugs are not as potent as traditional cut-offs (IC50 < 1um). For example, while 5-Fluorouracil and Doxorubicin are the backbone of many drug combinations, 5-Fluorouracil is often reaches clinical concentrations as high as 16,000 nM, while Doxorubicin is can reach concentrations closer to 19 nM – showing a 1000-fold difference in potency despite being two of the most commonly used oncology drugs.^30^ Another limitation is that typical drug assays are continual exposure to the drug for 24, 48, or 72 hours, while in *in vivo* systems, the drug is actively being cleared from the system. While our assay is a continuous-exposure model, we believe further work can improve clinical translation by incorporating PK-PD principles into cell-based assays. In the drug discovery sector, new machine learning and AI tools can predict C_max_, off-target toxicity, and other PK parameters.^31,32^

Additionally, many fields, such as antimicrobial research, focus on the minimum concentration required to inhibit growth, rather than just drug potency. There are now multiple methods for monitoring cell growth and disease progression over time using fluorescent microscopy in cancer.^12,13,33^ While these methods can provide key mechanistic insights or cell-to-cell variability, our goal was to develop a drug screening platform and data analysis pipeline to evaluate drug efficacy and translation. While Sorger’s Growth Rate model provides essential information for understanding a drug’s ability to inhibit cancer growth *in vitro*, Simeoni’s model allows for a more detailed understanding of drug kinetic selectivity that can be used in combination with concentration potency. Our goal is to provide computational tools and experimental methods to reframe how we think about oncology drug efficacy beyond just potency.

One limitation of our model is that it is restricted to one cell line per cancer type; further work in the lab aims to add more cell lines within each cancer type to resolve differences in proliferation rate and cellular resistance and better understand patients’ responses to specific chemotherapies. While continual efforts are made to improve the model, our study lays the foundation for analyzing complex drug responses to facilitate clinical translation and support either drug discovery or personalized methods. While the complexity of cell modeling systems needs to be improved, there is also a need to understand drug efficacy beyond traditional concentration-dependent metrics, incorporating pharmacodynamics and pharmacokinetics.

## Conclusion

There is a significant gap between *in vitro* assays and clinical translation that needs to be addressed to advance personalized medicine and improve the current drug development pipeline. Current drug discovery efforts are focused on concentration-dependent pharmacodynamic models. While these models are beneficial, they fail to account for the dynamic nature of drug response. Critically, both *drug concentration* and *exposure time* contribute to clinical efficacy, yet most laboratory studies focus on IC_50_ alone.^34^ We believe this indicates a need for a more holistic understanding of drug efficacy beyond traditional pharmacodynamic metrics like IC50, as well as for more complex models to account for the different factors contributing to drug response. Incorporating kinetic measures like minimum time for response (tmin), alongside Simeoni-style PK–PD modeling(Ct, ER, Kdmg, Kpro), provides a conceptual bridge from cell-based assays to real clinical outcomes. This framework highlights that potency alone is insufficient, and time-dependent killing must be measured to improve predictive power.

In this study, we applied the standard model for animal PK/PD studies in an *in vitro* assay, in combination with a novel continuous cell viability assay. This tool, along with the other methods outlined in this paper, enables researchers to process complex time-dose response curves using PK-PD and compare them with clinical response rates and indications. While these tools help bridge the gap in clinical translation by combining drug kinetics with traditional PD drug efficacy, they are not the only tools needed to advance the medicinal chemistry and personalized medicine fields. In future work, we plan to expand the cell lines and the drug library and to apply this method to complex cell models, such as 3D cultures and co-culture systems. We hope to reframe how we think about drug efficacy as both dose- and time-dependent to improve the rate of drugs reaching the market and to help patients receive the best oncology drugs for personalized medicine.

## Methods

### Cell Culture/Cell Lines

DU4475 (human breast cancer), DLD-1 (human colon cancer), and NALM6 (human B-ALL Leukemia) were obtained from ATCC and cultured according to ATCC recommendations. All were grown in RPMI 1640 with Glutamax (Gibco, Cat# 72400-047) and were fortified with 10% FBS (Fisher Scientific, Cat# FB12999102) and 1% Penicillin-Streptomycin (Gibco; Cat# 15140-122).

### Drugs

The drug classes chosen were the Anthracyclines (Doxorubicin, Epirubicin), Anti-metabolites (5-fluorouracil, Cytarabine), Topoisomerase 1 inhibitors (Topotecan, SN-38), Taxanes (Paclitaxel, Docetaxel), and the Vinca Alkaloids (Vincristine, Vinblastine). Drugs were obtained from SelleckChem as a 10 mM solution or reconstituted to a 10 mM solution in DMSO. The drugs were then diluted in a 10X dilution series in 1% DMSO in media for a final well concentration of 3.3 µM to 0.3 pM in 0.3% DMSO in Media. Eight different concentrations were used for each of the drugs. Two drugs were chosen from each standard drug regimen: Anthracyclines (Doxorubicin and Epirubicin), Anti-metabolites (5-Fluorouracil and Cytarabine), Topoisomerase 1 inhibitor (Topotecan and SN-38), Taxanes (Docetaxel and Paclitaxel), and Vinca Alkaloids (Vinblastine and Vincristine).

### Promega RTG

Cells were split 1-2 days before the RTG experiment. Each cell line was first tested for optimal density according to Promega’s RTG manual. The optimal densities determined for each cell line were: DU4475 – 8,000 cells/well, DLD-1 - 4,000 cells/well, NALM6 – 40,000 cells/well. The cells were counted using a hemocytometer prior to plating. The cells were plated in either suspension (DU4475 and NALM6) or adherent (DLD-1) white flat bottom Nunc 96 well plates (Thermo Scientific cat# 236105 for suspension and cat# 136101 for adherent). 0.3% DMSO in media and regular media controls were included on each plate. Each drug was tested in triplicate with 2 trials per drug (n=6). The drug dilutions (see Drugs subsection above, 100 µL/well) and 2X RTG reagents (100 µL/well, Promega, cat# G9712) were added after plating the cells (100 µL/well) for a total volume of 300 µL/well. A PBS border was included on the top and bottom row of the plate (300 µL) to reduce any potential plate effects throughout the experiment. The plate was incubated for 1 hour according to the Promega assay protocol. The RTG reagent concentration was calculated based on the Promega assay protocol. The plate was then transferred to the BioTek Synergy H1 plate reader which took luminescent measurements every 2 Hours for 72 hours. The plate reader was maintained at 37°C and 5.0% CO_2_ for the 72-hour time course. At the end of the 72 hours, the data was exported into an Excel file that was then used for the analyses.

### Data Analysis Pipeline

Each plate contained two of the drugs tested in triplicate along with controls. The raw RLU data were exported from the BioTek plate reader software, Gen5.14 (Fig. 2A). Each triplicate was averaged and compiled into a single Excel file containing drug, cell line, and trial number, which was imported into R-Studio for further analysis (Fig. 2A). The Viability Fraction, Growth Rate, and IC_50_ were all done in RStudio. The Average Relative Luminescence Units (RLU) for each replicate were compiled. The average viability fraction across trials was used to calculate the IC_50_ for each time point. The viability fraction was determined by dividing the luminescence of treated cells by that of untreated cells at each time point. This data was then used to calculate the IC_50_ using the Breeze drug screening scoring system, with code from the Breeze Pipeline.^35,36^ The fold change was determined by dividing the average luminescence by the luminescence at the initial time point. Sorger’s growth rate was determined by finding the slope, which was the fold change at the time point divided by the time lapsed for both untreated cells (k_CTRL_) and treated cells (k_DRUG_) and using the Sorger Growth Rate Equation outlined in Fig. 1B.^21,22^ The data from the fold change, viability fraction, growth rate, and IC_50_ were visualized using ggplots and heatmap functions in R-Studio.. The color gradient was assessed for colorblind accessibility as a light (red) to dark (green) for deuteranopia (6%) using the ColorBlindness package in R-Studio. The data were saved to an external file for data visualization with the RShiny package. The raw data and code used to generate the figures can be found at https://github.com/fingolfn/rtg_simeoni.

The Simeoni Tumor Growth Inhibition Model was performed using the method outlined in Simeoni Methods Section and Monolix Suite (https://lixoft.com/) Software from the Average RLU Divided by the Initial RLU values. The parameters were exported from Monolix and used for downstream analysis, such as Concentration Threshold in R-Studio. The concentration threshold was determined using the Simeoni reformulation method below (Eq. 09-12). The parameters determined from the fit were then used to create predicted models, which were compared with the raw data to assess the fit’s accuracy (Fig. 4). While each drug for a particular cell line was run simultaneously, the parameters were not assumed to be dependent on the concentration tested. Originally, the [C_pro_] parameter was set to 1 under Sorger’s idea of comparing to “initial cell density”, but this led to peaks occurring later than the data indicated.

### PCA Analysis and ORR Prediction Model

Principal Components Analysis (PCA) was conducted on the viability fraction and growth rate datasets using the prcomp function, and the results were visualized in RStudio using plotly. The responding data, in which the growth rate or viability fraction dropped below 0.5, were then used to determine the driving factors in the pharmacodynamic response by applying the same prcomp function (Supp Fig 1). To model how the efficacy ratio and the IC_50_/C_max_ Ratio relate to clinical ORR, a generalized additive model (GAM) was fit using the mgcv package in RStudio. A beta regression with a logit link was applied to predict ORR, as the fraction of patients responding is bound between 0 and 1. To account for the potential nonlinear relationship between the parameters and ORR, a smoothing function was applied to the model [s(IC_50_/C_max_, ER)]. Predicted ORR values were generated over a range (-4 to 4) and visualized alongside the experimental values using ggplot2.

### Simeoni reformulation

#### Original form of governing equations

Building on the Simeoni PK-PD framework, we re-expressed the cell-fate compartments to make each kinetic constant interpretable in standard pharmacological terms. Tumor burden is represented as concentrations of proliferating ([C]pro) and damaged ([C]_dmg_) cells per unit volume (L^−1^). Pure exponential growth (see details in supporting information) is assumed because the expressions for the time-efficacy index (TEI) and threshold concentration (Ct) arise analytically only in this regime. The proliferation rate constant is denoted kpro (s^−1^). Drug action is split into (i) a toxic conversion step with rate k_kill_ ≡ k1 (s^−1^), and (ii) an irreversible damage propagation step with rate k_dmg_ ≡ k2 (M^−1^ s^−1^). Because k_dmg_ is second-order in units, it subsumes classical potency metrics such as the IC_50_;(M^−1^ s^−1^) higher potency lowers the drug concentration required to achieve a given damage flux. This fact that k_2_/k_dmg_ is an aggregate parameter is also mentioned explicitly in the original Simenoni paper where: “k2, is a measure of drug potency” is stated five times. Under a single-step approximation (no transit compartments), the system:

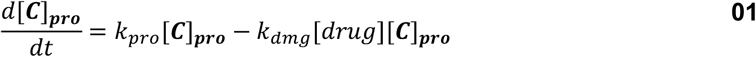

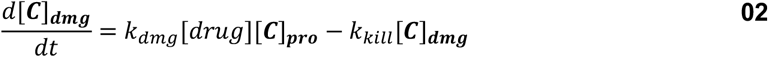

admits closed-form solutions that map directly onto the original Simeoni TEI and Ct expressions while clarifying the physiological meaning of each parameter:

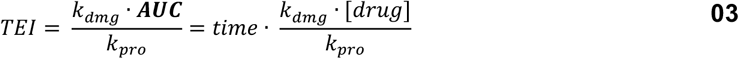

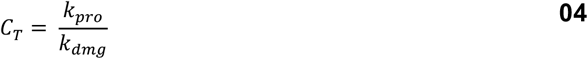

#### Incorporation of a Hill-type drug–response relationship

To enable direct comparison between kinetic efficacy metrics and conventional potency measurements, we replaced the linear drug term in the damage equation with a Hill function that partitions the cell population into actively proliferating and drug-affected fractions:

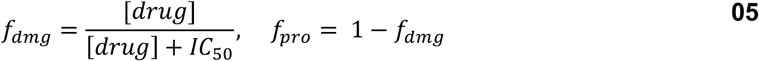

Here IC_50_ is the concentration giving half-maximal conversion of proliferating to damaged cells. Substituting f_pro_×[C]_total_ and f_dmg_×[C]_total_ for [C]_pro_ and [C]_dmg_ in the differential system yields analytic forms for C_t_ and TEI that now include an explicit IC_50_ term. This linkage allows experimental IC_50_ values to be benchmarked against kinetic surrogates of efficacy without additional fitting.

#### Analytical solutions for efficacy metrics

Assuming drug exposure is non-saturating ([Drug] ≪ IC_50_) the model simplifies to two limiting cases that again reduce to pure exponentials. In both limits, we recover Ct and TEI:

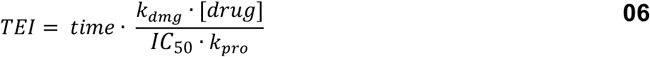

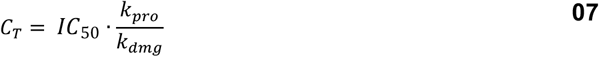

These closed-form results make it possible to translate dose-response data into time-based efficacy projections and to compare compounds on a common mechanistic scale that integrates proliferation dynamics, potency, and pharmacokinetics. Critically, under this formulation both rate constants are first order which yields a unitless ratio we call an efficacy ratio:

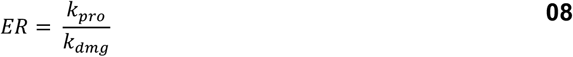

We use this ratio to capture the kinetic “competition” of proliferation vs damaging rate constants where ER > 1 reflects net proliferation while ER < 1 reflects net cytotoxicity.

The original Simeoni Equations were reformatted with the new terms for the equations above (Eq. 01-08):

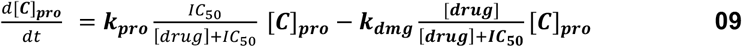

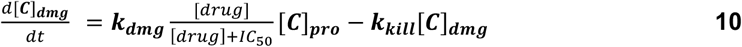

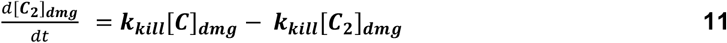

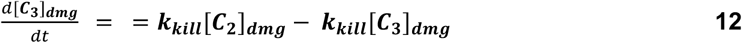

### Regimen Calculations (C_max_)

The maximum plasma concentration (C_max_) was determined by compiling the different drug administration regimens based on Cancer Principles and Practice of Oncology.^20^ The C_max_ was calculated using typical PK parameters for the drug based on Obach, et al.^34^ The Average C_max_ was determined by taking the geometric mean of the highest C_max_ and the lowest C_max_ based on the different dosing regimens and schedules that could be given for any particular dose.

### ORR Data Compilation

Our single-agent dataset (Fig. 3A) compiles overall response rates (ORR) of first-line treatment of metastatic cancer therapies from phase 2 trials published between 1950 and 2000. To build a chemotherapy response-rate dataset, we first encoded raw data published by the National Cancer Institute (NCI) as a textbook in 1970.^37^ This dataset (“ORR Matrix”) included data on 19 currently indicated therapies from 786 clinical trials and 19,958 evaluable patients across 19 cancer types. This dataset was updated with meta-reviews published between 1970 and 2000 to fill gaps and add 25 drugs (“NCI1970meta”) producing single-agent response dataset across 49,002 evaluable patients (Fig, 3A). The result was an analysis of 30 drugs in 17 different cancer lineages.^24^

