## Supplemental Figures for "Development of a Time-based Drug Screening Platform for Improved Clinical Translation"

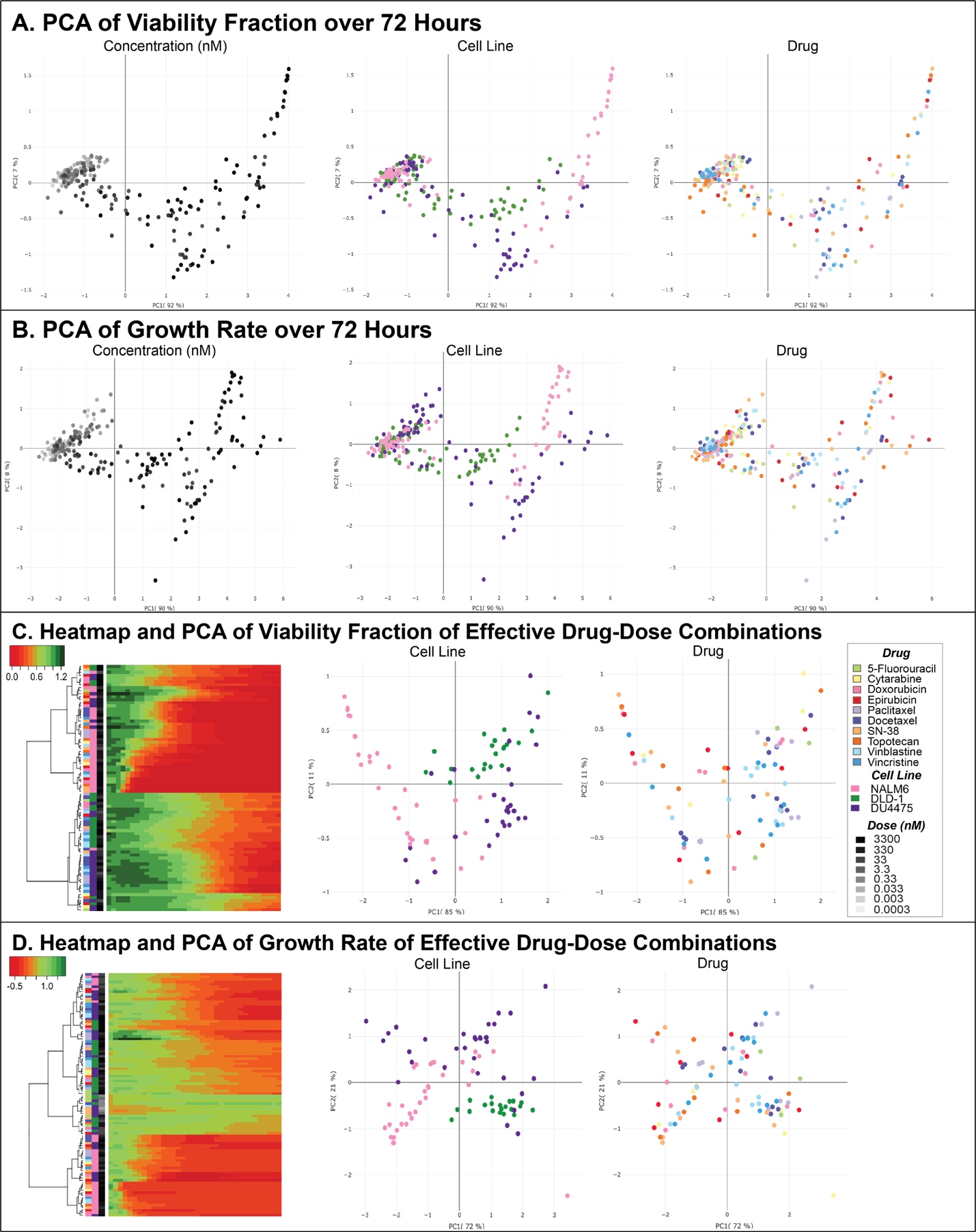

**Supplemental Figure 1 – Analysis of Trends in Viability Fraction and Growth Rate Data Using Principle Component Analysis.** Color key for experimental parameters can be found on the right side of panel C. **A-B.** Principals Components Analysis (PCA) on the Viability Fraction Matrix (**A**) and Growth Rate Matrix (**B**) from Fig. 2A with different experimental parameters colored for each graph: concentration (left), cell line (middle) and drug (right). **C-D.** Truncated matrix of dose-cell line-drug combinations that had a minimum of a 50% response for Viability Fraction (**C**) and Growth Rate (**D**) was used for PCA to determine the driving factor of drug sensitivity cell line (right) or drug (right).

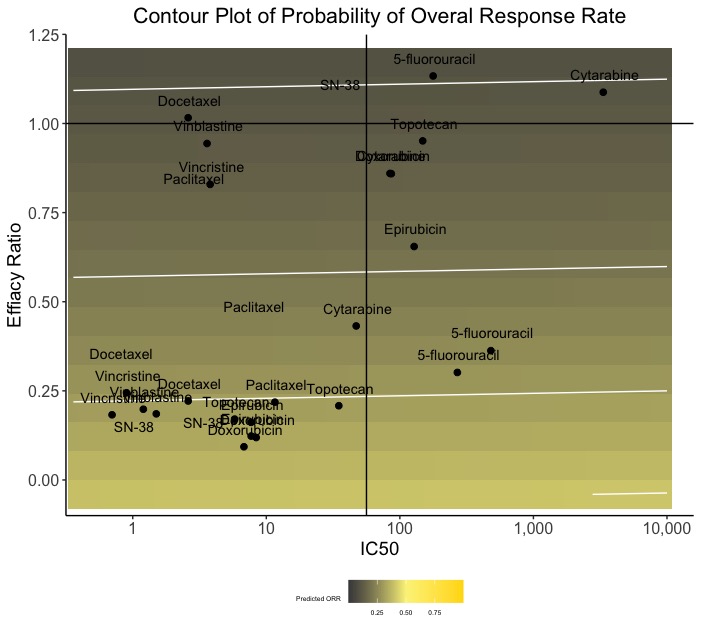

**Supplemental Figure 2: Prediction of ORR using Efficacy Ratio and IC50. .** A contour plot of probability of patient response rate developed using a generalized additive model using the data from Fig. 2D and Fig. 4C

**Supplementary Table 1. Values of The Simeoni-TGI Model Applied *In Vitro***

The values used for analysis and graphics from the Simeoni-TGI Model in our experimental data

| **Drug** | **Cell Line** | **K_PRO_**  **(Max) (s^-1^)** | **K_DMG_ (Max) (nM/s)** | **K_KiLL_**  **(Max) (s^-1^)** | **Efficacy Ratio** | **ORR** | **IC50 (nM) at 73Hr** | **Ct (nM)** |
| --- | --- | --- | --- | --- | --- | --- | --- | --- |
| 5-fluorouracil | DLD-1 | 0.06 | 0.06 | 0.05 | 0.94 | 0.22 | 266.50 | 249.86 |
| 5-fluorouracil | DU4475 | 0.03 | 0.09 | 0.09 | 0.28 | 0.26 | 404.60 | 114.95 |
| 5-fluorouracil | NALM6 | 0.03 | 0.15 | 0.48 | 0.21 | 0.05 | 720.90 | 152.23 |
| Cytarabine | DLD-1 | 0.06 | 0.05 | 0.04 | 1.04 | 0.09 | 5000.00 | 5219.65 |
| Cytarabine | DU4475 | 0.03 | 0.08 | 0.15 | 0.36 | 0.09 | 126.50 | 45.37 |
| Cytarabine | NALM6 | 0.03 | 0.10 | 1.26 | 0.32 | 0.27 | 70.80 | 22.52 |
| Docetaxel | DLD-1 | 0.06 | 0.05 | 0.06 | 1.15 | 0.00 | 3.90 | 4.47 |
| Docetaxel | DU4475 | 0.01 | 0.05 | 0.23 | 0.23 | 0.59 | 3.80 | 0.86 |
| Docetaxel | NALM6 | 0.03 | 0.10 | 0.87 | 0.26 | NA | 1.30 | 0.33 |
| Doxorubicin | DLD-1 | 0.06 | 0.11 | 0.06 | 0.52 | 0.09 | 129.60 | 67.88 |
| Doxorubicin | DU4475 | 0.01 | 0.04 | 0.54 | 0.34 | 0.37 | 12.50 | 4.30 |
| Doxorubicin | NALM6 | 0.04 | 0.69 | 1.03 | 0.06 | 0.58 | 10.20 | 0.58 |
| Epirubicin | DLD-1 | 0.03 | 0.01 | 0.05 | 3.66 | 0.07 | 192.00 | 703.43 |
| Epirubicin | DU4475 | 0.01 | 0.02 | 0.58 | 0.37 | 0.31 | 11.50 | 4.27 |
| Epirubicin | NALM6 | 0.04 | 0.43 | 1.30 | 0.09 | 0.74 | 11.60 | 1.02 |
| Paclitaxel | DLD-1 | 0.05 | 0.07 | 0.06 | 0.66 | NA | 4.20 | 2.77 |
| Paclitaxel | DU4475 | 0.01 | 0.08 | 0.21 | 0.13 | 0.33 | 17.50 | 2.24 |
| Paclitaxel | NALM6 | 0.03 | 0.08 | 0.72 | 0.34 | NA | 11.90 | 4.05 |
| SN-38 | DLD-1 | 0.05 | 0.05 | 0.08 | 0.96 | NA | 52.60 | 50.63 |
| SN-38 | DU4475 | 0.01 | 0.12 | 0.38 | 0.10 | NA | 4.90 | 0.51 |
| SN-38 | NALM6 | 0.03 | 0.39 | 2.44 | 0.09 | NA | 1.50 | 0.13 |
| Topotecan | DLD-1 | 0.05 | 0.11 | 0.06 | 0.47 | 0.10 | 222.70 | 105.61 |
| Topotecan | DU4475 | 0.01 | 0.06 | 0.39 | 0.20 | 0.15 | 52.50 | 10.34 |
| Topotecan | NALM6 | 0.03 | 0.36 | 2.04 | 0.09 | 0.00 | 8.70 | 0.75 |
| Vinblastine | DLD-1 | 0.04 | 0.04 | 0.07 | 1.16 | 0.07 | 5.40 | 6.27 |
| Vinblastine | DU4475 | 0.01 | 0.05 | 0.18 | 0.24 | 0.20 | 2.30 | 0.54 |
| Vinblastine | NALM6 | 0.03 | 0.22 | 1.14 | 0.16 | 0.25 | 1.80 | 0.28 |
| Vincristine | DLD-1 | 0.05 | 0.06 | 0.05 | 0.88 | 0.00 | 5.70 | 5.01 |
| Vincristine | DU4475 | 0.01 | 0.07 | 0.21 | 0.15 | 0.21 | 1.10 | 0.16 |
| Vincristine | NALM6 | 0.04 | 0.27 | 1.10 | 0.13 | 0.52 | 1.30 | 0.17 |

**Supplementary Table 2. Pharmacokinetic Parameters in nM**

The maximum plasma concentration (C_max_) was determined by compiling the different drug administration regimens based on Cancer Principles and Practice of Oncology.^18^

| **Drug** | **C_max_ Range – low** | **C_max_ Range - high** | **Cmax Geometric Mean** | **TR_min_** | **TR_max_** |
| --- | --- | --- | --- | --- | --- |
| **5-fluorouracil** | 760 | 320,000 | 15,595 | 3460 | 4610 |
| **Cytarabine** | 130 | 49,000 | 2524 | 180 | 1800 |
| **Docetaxel** | 14 | 56 | 28 | - | - |
| **Doxorubicin** | 9.8 | 35 | 19 | 3.1 | 10 |
| **Epirubicin** | 13 | 25 | 18 | 4.6 | 23 |
| **Paclitaxel** | 50 | 240 | 110 | 14 | 140 |
| **Irinotecan** | 340 | 7700 | 1618 | 850 | 17,000 |
| **Topotecan** | 1.5 | 27 | 6.4 | 1.5 | 27 |
| **Vinblastine** | 0.54 | 0.92 | 0.7 | - | - |
| **Vincristine** | 2.6 | 7 | 4.3 | - | - |

**Supplementary Table 3. Simeoni-TGI Parameters from Original Publication**^20^

The literature and experimental values reported by Simeoni and colleagues for clinical validation of their model.

| **Drug** | **Clearance (L/m^2^/Hr)** | **k_PRO_**  **(Hr^-1^)** | **k_DMG_**  **(nM/Hr)** | **Systematic Exposure (nM*Hr)** | **Mid Clinical Dose (nM/m^2^)** | **ER** | **Ct (nM)** |
| --- | --- | --- | --- | --- | --- | --- | --- |
| **5-fluorouracil** | 39.5 | 0.01 | 0.01 | 608.20 | 24024.05 | 0.0009 | 0.92 |
| **Docetaxel** | 22.6 | 0.01 | 0.22 | 4.79 | 108.31 | 0.0001 | 0.06 |
| **Doxorubicin** | 40 | 0.02 | 0.37 | 3.22 | 128.79 | 0.0000 | 0.04 |
| **Irinotecan** | 14.8 | 0.01 | 0.01 | 51.66 | 764.52 | 0.0004 | 0.44 |
| **Paclitaxel** | 13.6 | 0.01 | 0.02 | 18.30 | 248.86 | 0.0005 | 0.54 |
| **Vinblastine** | 29 | 0.01 | 1.28 | 0.43 | 12.33 | 0.0000 | 0.01 |
| **Vincristine** | 4.92 | 0.01 | 3.79 | 0.37 | 1.82 | 0.0000 | 0.00 |
| **Cisplatin** | 15.6 | 0.02 | 0.08 | 17.03 | 265.69 | 0.0002 | 0.19 |
| **Etoposide** | 1.68 | 0.01 | 0.01 | 333.75 | 560.69 | 0.0028 | 2.80 |
| **Gemcitabine** | 91.8 | 0.01 | 0.03 | 118.99 | 10923.34 | 0.0005 | 0.49 |
